# The hand microbiome is sensitive to topical antibiotics and has varying sensitivity to liquid soaps

**DOI:** 10.64898/2026.06.30.735469

**Authors:** M. Stenton, S.R. Henderson

## Abstract

Hand eczema has been described as having an increased prevalence in persons with increased frequency of hand washing. This study investigated the differences in the hand microbiome of persons with and without a history of eczema and secondly the sensitivity of these microbes to commercial liquid soap as a potential trigger for eczema flares. The study identified Staphylococcus to be the most populus genus on the hands in both groups, but the distribution of species was different. Additionally, there was no difference in the number of soaps that produced zones of inhibition but there were some differences in the overall sensitivity to the different soaps tested. Overall, it was determined that liquid soap can cause bactericidal effects on some species of the commensal microbiome, but further work is required to determine if this could be the cause of hand eczema.

## Introduction

Eczema, a common inflammatory skin condition, with a primary on set in early childhood is characterised through patches or lesions of skin with an abnormal skin barrier resulting in changed appearance and are often described as being “itchy”. These lesions can flare or worsen in severity or can improve and go into remission with appropriate management (Girolomoni & Busà, 2022). Atopic dermatitis (AD) is the predominant cause of eczema and eczema like skin conditions. During lesions occurring within AD it has been repeatedly observed that there is typically a dysbiosis in the skin’s microbiome, with increased severity of AD being correlated with a low Shannon index of microbial diversity alongside an unusual abundance of *Staphylococcus aureus* on the skin (Leyden, Marples & Kligman, 1974; Khadka *et al*., 2021), therefore it is critical for us to study in depth the basic factors that might be causing this decrease in the diversity within the skin microbiome in AD and other conditions such as hand eczema. It has been observed that there is a correlation between persons in positions that require high frequency handwashing, what remains unclear is whether this is due to disruption of the microbiome or direct impact on skin inflammation (Lauharanta *et al*., 1991; Ibler, Jemec & Agner, 2012).

The specific cause of eczema has not been determined hence there is a possibility that the causative agent of hand eczema is via the healthy microbiome being depleted by the antimicrobial agents in antimicrobial soap. Therefore, this study investigated if strains isolated from persons with a history of eczema had a higher degree of sensitivity than those isolated from persons with no history of eczema. In this study strains of bacteria were obtained from healthy individuals without inflammatory diseases who have either had eczema as a child or never had eczema. A range of commercially available soaps were then tested against the bacterial strains to determine if antimicrobial soaps decreased the growth of bacterial strains obtained from persons with a history of eczema to a greater extent than those without a history of eczema.

## Materials and Methods

### Microbiome collection

PreMoise Hygiene Swab (Technical Service Consultants Ltd) were rubbed repeatedly over the upper (dry) surface of the hand and fingers to collect the microbiome of consulting adults without inflammatory co-morbidities. These were declared to have either never had eczema (Healthy) or to have previously had periods of eczema (Eczema) on consent. The swabs were spread over Columbia 5% defibrinated horse blood agar (CBA) or Tryptic Soy 5% defibrinated horse blood agar and Mannitol salt phenol red agar (MSA) plates and incubated at 37°C for 2 days. Individual colonies (8-10) were selected and grown in Brain heart infusion broth or Nutrient broth to stationary phase and frozen with 25% glycerol.

### Speciation

Glycerol stocks were streaked on CBA plates and grown to achieve single colonies. Single colonies were picked into 30 µl Tris-EDTA buffer and incubated at 95°C for 10 minutes. Remaining colonies were pelleted at 6000 xg for 10 minutes. 2 µl supernatant was used as the template for 25 µl colony PCR reactions in 1x DreamTaq DNA Polymerase mastermix (ThermoFisher) with the 8F and 907R primers (Table 1). Thermocycling (95°C 4 minutes; 30 cycles of 95°C 30 s 50°C 30 s, 72°C 60 s; 8 minute final extension at 72°C) was performed, amplification confirmed with agarose gel electrophoresis. PCR products were purified using a QIAquick PCR Purification Kit (Qiagen) and sequenced from the 27F primer (AGAGTTTGATCATGGCTCA) (Eurofins Genomics). Obtained sequences were compared to the 16S ribosomal RNA sequences database and confirmed to genus level (NIH). Staphylococcal strains were confirmed where ambiguity existed utilising a 25 µl multiplex colony PCR reaction with the epi-F/R, cap-F/R, sap-F/R and rae-F/R pairs of oligonucleotides in 1x DreamTaq DNA polymerase mastermix. Thermocycling was performed with 95°C 5 minutes; 30 cycles of 95°C 30 s, 58°C 30 s, 72°C 70s; 72°C 2 minutes. Fragments were analysed by agarose gel electrophoresis (Table 1)(Hirotaki *et al*., 2011).

**Table 1:**
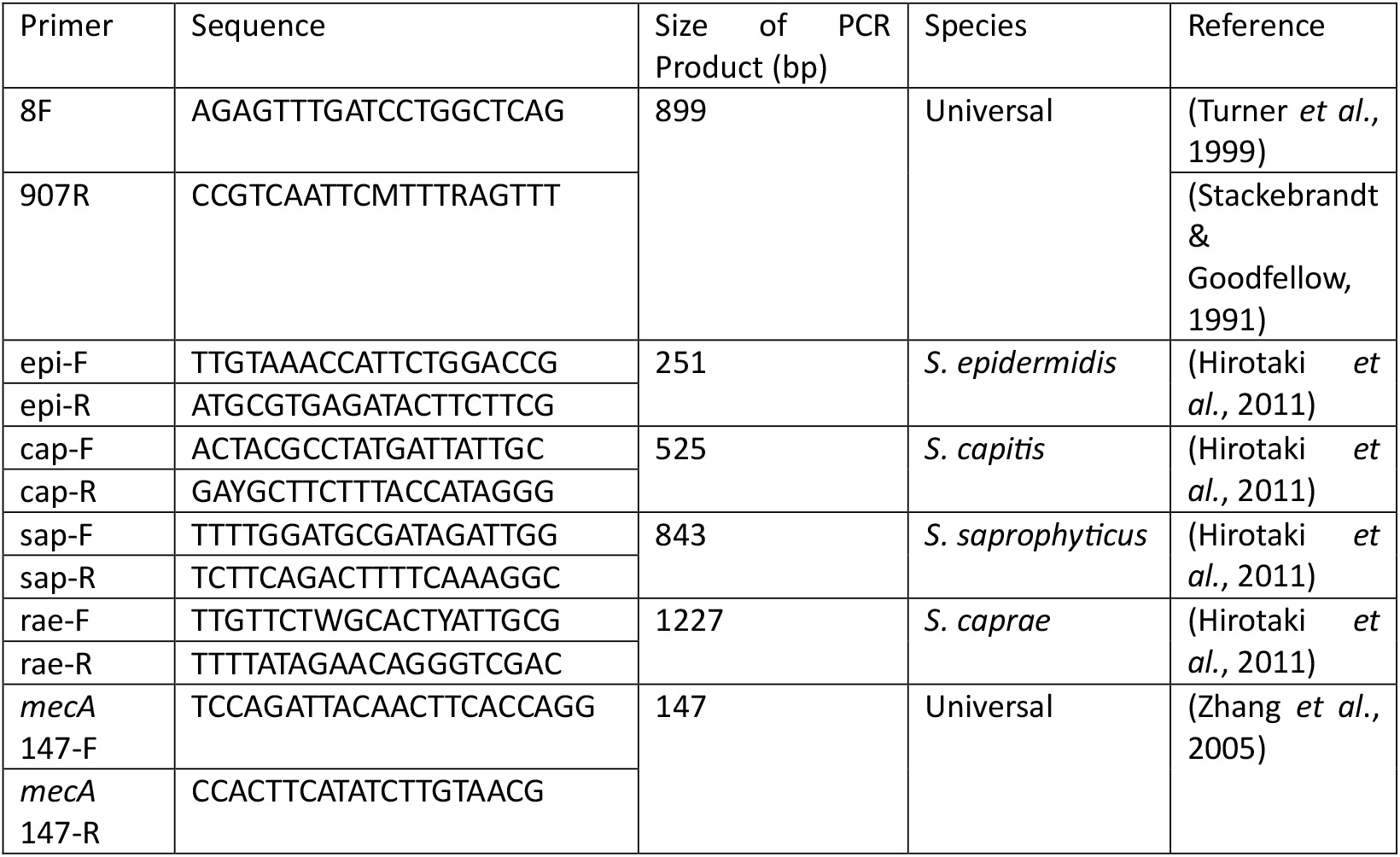
Oligonucleotide primer sequences for PCR amplification.

### *mecA* resistance testing

Staphylococcal species were tested for the presence of the SCC*mec* through PCR screening for the *mecA* gene using the *mecA* 147 primers as per the conditions of (Zhang *et al*., 2005).

### Sensitivity testing

Kirby-Bauer disk diffusion assays were performed on Mueller-Hinton agar. A lawn was made from a bacterial resuspension at OD600 = 0.1. Twenty-one commonly available liquid soaps were bought from Amazon. Two contained chlorhexidine gluconate (J,L) and a further 11 had antibacterial claims (B,C,D,F,G,M,N,O,R,S,U). Soap (20 µl) was loaded onto Oxoid blank antimicrobial susceptibility disks and placed onto the agar. Neomycin (10 µg) and fusidic acid (10 µg) Oxoid disks were used as controls. Plates were incubated at 37°C for 18 hours or until confluent.

## Results and Discussion

### The diversity of the microbiome on the hands

The microbiome of the dry skin of the human hands from healthy individuals with and without a history of eczema was sampled and a selection (maximum 8 per sampling event) of isolated colonies were retained for additional analysis. Four of the colonies were found to be unculturable under the trialled conditions and not retained for further analysis. The Staphylococcus genus was found to be the most prevalent genus from the samples analysed accounting for 58% of the total strains collected (50% of strains from persons with an Eczema history and a total of 62% of those without) (Figure 1). Only one of the culturable strains was not identified (Eczema) due to failed PCR reactions. Acinetobacter and Pantoea were found only in healthy individuals whereas Moraxella was found only in individuals with an Eczema history. However, as all of these are at low frequency it cannot be ruled out that they are a very low frequency and were not detected.

**Figure 1:**
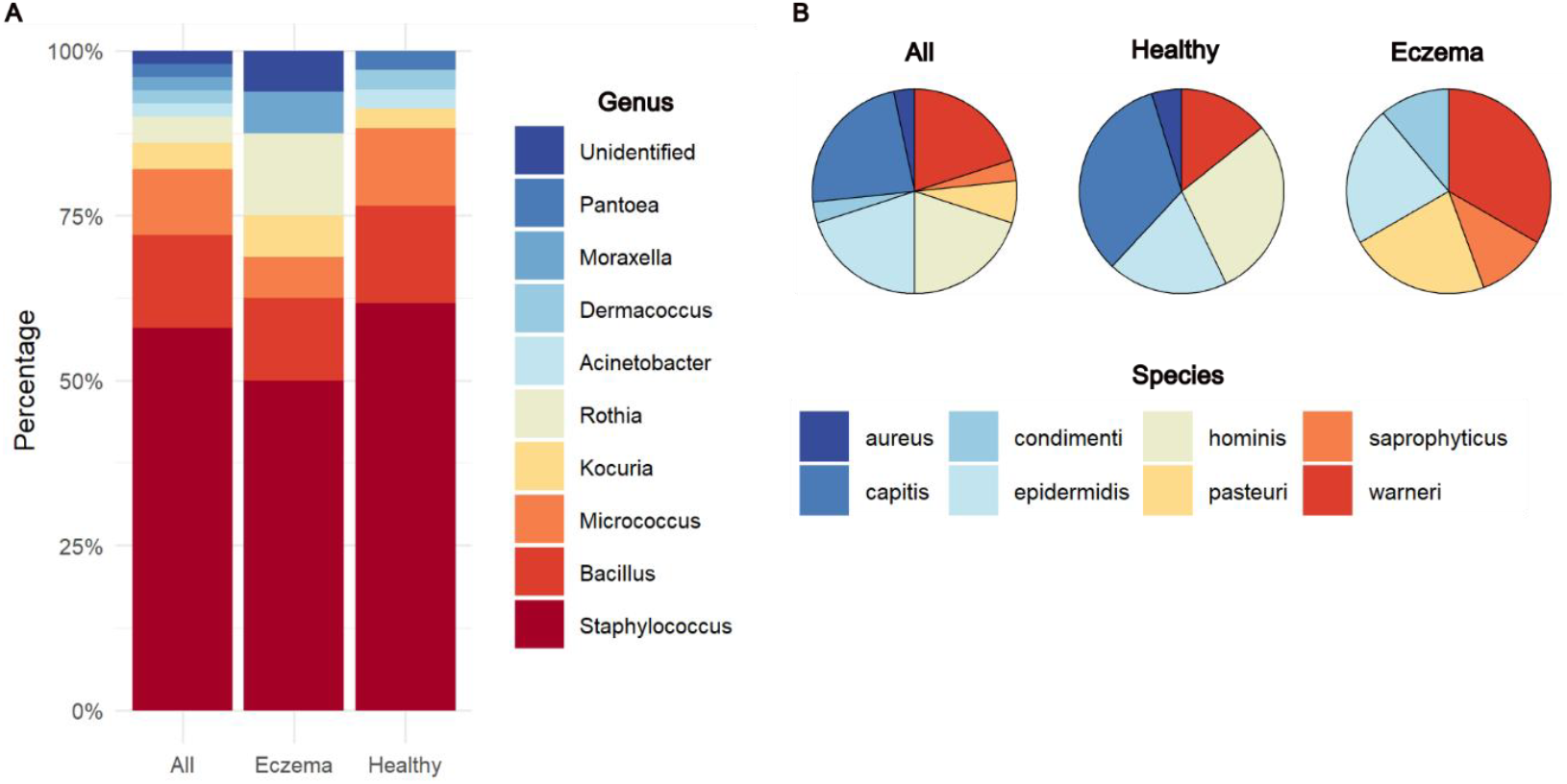
The diversity of genus of bacteria (A) or species of Staphylococci (B) isolated from all samples (All), healthy persons with a history of Eczema (Eczema) and healthy individuals without a history of eczema (Healthy) adjusted as a percentage of the total isolated strains. Genus and species were identified using 16S rRNA amplification and sequencing with multiplex Staphylococcus PCR where required.

It was possible to perform subspeciation of the Staphylococcus genus through use of 16S rRNA sequencing and additional multiplex PCR analysis to speciate *Staphylococcus capitis* and *Staphylococcus caprae* which share a common 16S rRNA sequence (Hirotaki *et al*., 2011) (Figure 1). Previously, *Staphylococcus aureus* has been described as a transient coloniser of healthy skin, but with being a high-frequency coloniser of atopic dermatitis lesions (Leyden *et al*., 1974). In this study only a single isolate of *S. aureus* was isolated, and more interestingly, no isolates of *S. aureus* were isolated from persons with a history of eczema. The second major observation being that in contrast to the stability of the genus level identifications being relatively stable between the two groups. There was a relative shift in the diversity in staphylococci identified in the two different groups. In healthy individuals the most frequent species were the capitis/caprae and hominis which were not identified on any person who had previously had eczema. In comparison, pasteuri and saprophyticus were uniquely identified in persons with a history of eczema but not on persons with continuous health. The only staphylococci determined in both groups were epidermidis and warneri which was the most common in persons with history of eczema but less frequent in the healthy group. Previous studies of the hand microbiome have preferentially focussed on the moist palm surface of the hands and the finger tips, as opposed to the dry dorsal surface of the hands and the fingers which is primarily impacted by hand eczema (Edmonds-Wilson *et al*., 2015). The density of *S. hominis* and *S. epidermidis* were previously determined not to be influenced by the presence of hand eczema and that *S. epidermidis* was the most prevalent Staphylococci on the hands of persons without hand eczema (Nørreslet *et al*., 2022), however in our study we did not identify any *S. hominis* on the hands of persons with a history of eczema, and the most prevalent Staphylococcal species was found to be *S. capitis* in the always heathy group. A limiting factor of our study was only a relatively small number of individual strains were selected for further study and hence a full analysis of the diversity is not possible from these results.

### Methicillin Resistance

After confirmation of the bacterial genus, the strains which were defined as staphyloccocal were further characterized to determine the extent of which the SCC*mec* mobile genetic element was present within the sample set. Of the 21 samples tested from persons with a healthy background 3 gave a positive PCR result for containing the *mecA* gene (supplementary figure 2). On the other hand, none of the 9 strains tested from persons with a history of eczema gave a positive PCR result for *mecA*. Previous findings have shown that between 0-18.3% of Atopic dermatitis patents colonised with *S. aureus* carried methicillin resistant *S. aureus* (Chung *et al*., 2008; Suh *et al*., 2008; Balma-Mena *et al*., 2011; Petry *et al*., 2014) correlating with a 16% MRSA nasal colonisation as found by (Davis *et al*., 2004). While our study focuses on the coagulase negative Staphylococci of the commensal skin microbiome which has previously been found to have a 13.5% incidence of oxacillin resistance (Garbacz *et al*., 2021). Our carriage rate of 14.2% within the healthy background group is in line with this figure. However, surprisingly the group with historical eczema who were likely to have enhanced topical antibiotic usage were not found to have any incidence of *mecA* carriage. One explanation is that there is greater variability in the SCC*mec* in the coagulase-negative species range, with increased incidence of *mecC* which the PCR screen did not account for.

### Comparison of Soap Sensitivity

A sub-selection of the isolated strains was further tested for the sensitivity to 20 commercially available liquid soaps and two topical antibiotics (neomycin and fusidic acid) (Figure 2). Both neomycin and fusidic acid inhibited the growth of all strains with the zones of inhibition all being greater than 10 mm. As many of the strains are primarily considered to be commensal organisms the zones of inhibition were not compared against reference tables. Two stocks were found to have mixed populations [*S. capitis* and *Micrococcus spp*.] and [*S. hominis* and *S. capitis*] and not tested for sensitivity, and one stock of *S. warneri* was found not to be viable in liquid culture.

**Figure 2:**
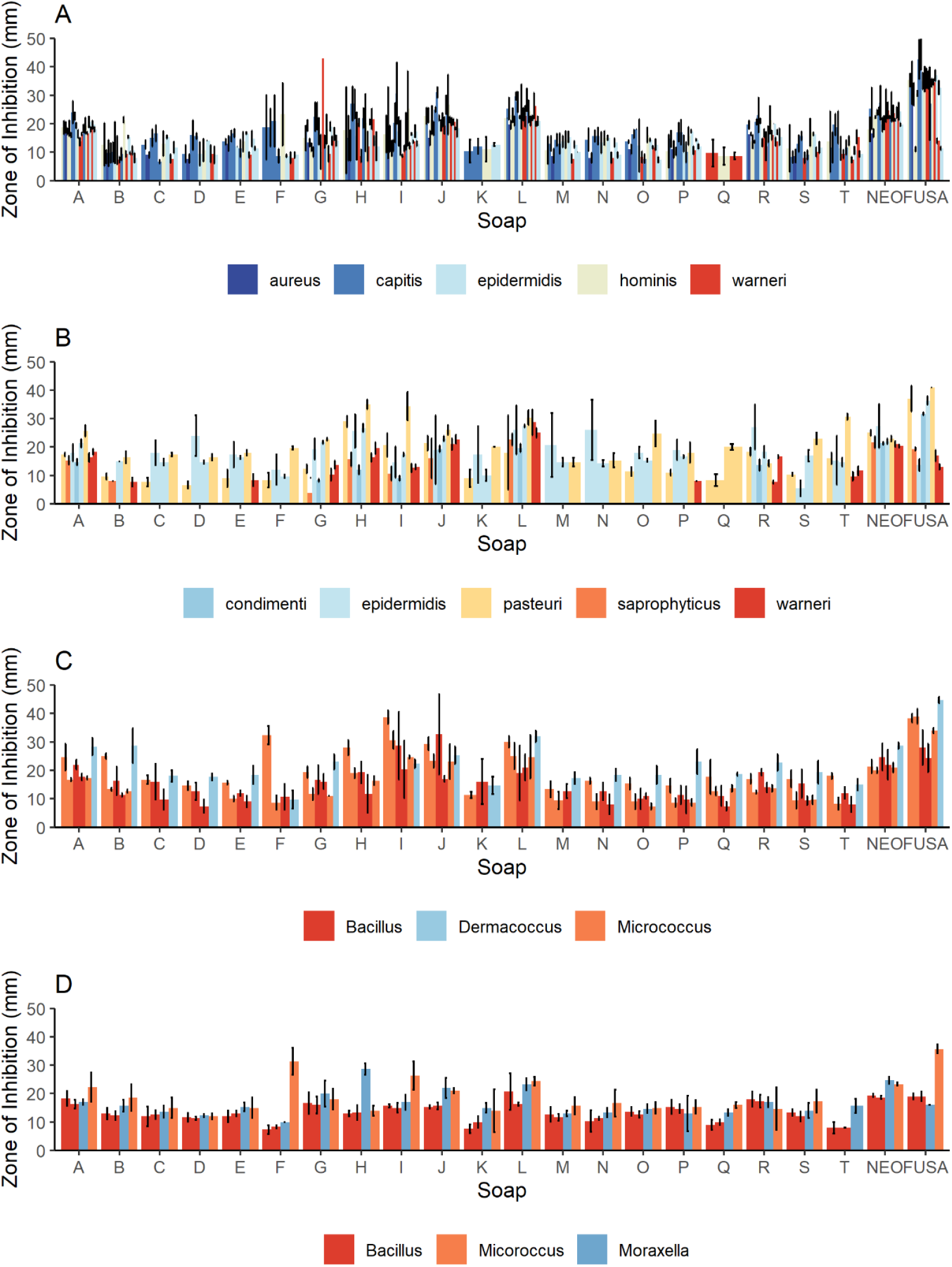
The zones of inhibition measured in millimetres obtained from soaps A-U and Neomycin (NEO) or Fusidic Acid (FUSA). (A) Staphylococcal strains isolated from healthy individuals with no history of Eczema. (B) Staphylococcal strains isolated from healthy individuals with a history of Eczema. (C) Non-staphylococcal strains isolated from healthy individuals with no history of Eczema. (D) Non-staphylococcal strains isolated from healthy individuals with a history of Eczema.

Of note one strain from the non-Staphylococcal group was subsequently after testing found to be a mixed isolate containing both *Staphylococcus capitis* and *Micrococcus spp*. (aloeverae/luteus) and as such was only included in the non-Staphylococcal group.

In general, a loss of zone of inhibition indicates resistance to an agent, therefore a general analysis of the overall sensitivity (soaps producing a clear zone of inhibition) was analysed. 2-tailed Student’s T-tests were performed and determined that there was a significant difference between in the percentage of strains with resistance to the tested soaps with the overall comparison of all strains from healthy individuals and those with a history of eczema having no comparable difference in their resistance to the tested soaps (P=0.33). There was no difference found within the staphylococcal strains between the two groups (P=0.84) but a significant increase in resistance was found in the non-staphylococcal strains (P=0.006) (Table 1).

Soaps J and L contained chlorhexidine gluconate at 1% (w/v) and 4% (w/v) respectively and were specifically marketed towards persons with eczema. All the commensal strains isolated were sensitive to these strains with zones of inhibition between 10 and 40 mm resulting from these soaps. Previous studies into the antimicrobial properties of chlorhexidine gluconate have given rise to decreased recovery of the skin microbiome in regions of treatment (Adams *et al*., 2005; Carty *et al*., 2014). While decreased sensitivity to a range of bacterium has been described, complete failure to prevent growth has not (Wang *et al*., 2008; Sheng *et al*., 2009; Hughes & Ferguson, 2017). This is in line with our results which indicate that while the commensal microbes tested had a range of susceptibilities complete resistance was not observed.

Two further soaps (A and I) also produced zones of inhibition against all tested strains despite neither carrying branding to indicate them to be antimicrobial. The two soaps with the least antibiotic action were K and Q which resulted in zones of inhibition in 16 of the 38 strains tested. Of the soaps marketed to be antimicrobial only three resulted in zones of inhibition to more than 80% of the tested strains (B, G, and R) (Table 1).

**Table 1:**
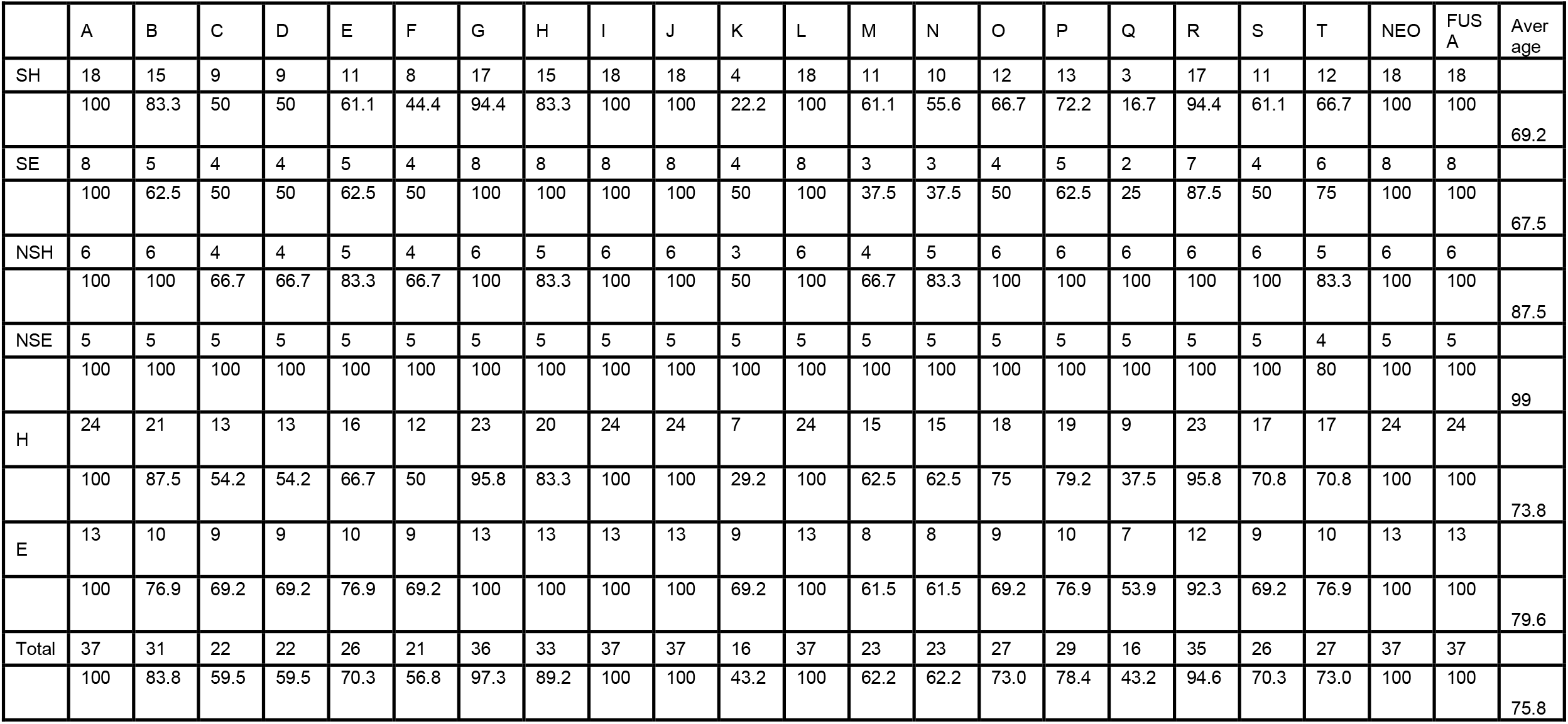
Number of strains and percentage of strains tested that resulted in a zone of inhibition to each soap and the control antibiotics. Soaps A-U and neomycin (NEO) and Fusidic Acid (FUSA). SH: Staphylococcus strains isolated from healthy individuals; SE: Staphylococcus strains isolated from healthy individuals with a history of eczema; NSH: Non-Staphylococcus strains isolated from healthy individuals; NSE: Non-Staphylococcus strains isolated from healthy individuals with a history of eczema; H: All strains isolated from healthy individuals; E: All strains isolated from healthy individuals with a history of eczema; Total: All strains isolated from all individuals.

Previous studies have investigated the long-term effects of antimicrobial soaps and disinfectant use on the hand microbiome, with differential conclusions. (Fierer *et al*., 2008; Two *et al*., 2016; Kramer *et al*., 2024) all concluded that while the proportions of bacteria were altered no lasting effect on the diversity was observed, whereas in a naïve population a lasting effect was observed (Yu *et al*., 2018). Our study suggests that commensal bacteria have differential sensitivities to liquid hand soaps and therefore this may account for transient suppression of the bacterial microbiome of organisms with the greatest sensitivity at the time of the hand wash, but with the ability to redistribute in the periods between the washing events.

## Acknowledgements

The work was supported by a National Eczema Association Engagement grant NEA23-196. Work was done in accordance with University of Bradford ethics committee EC28120.

## Data Availability

N/A

## Supplementary Figures

**Supplementary Figure 1:**
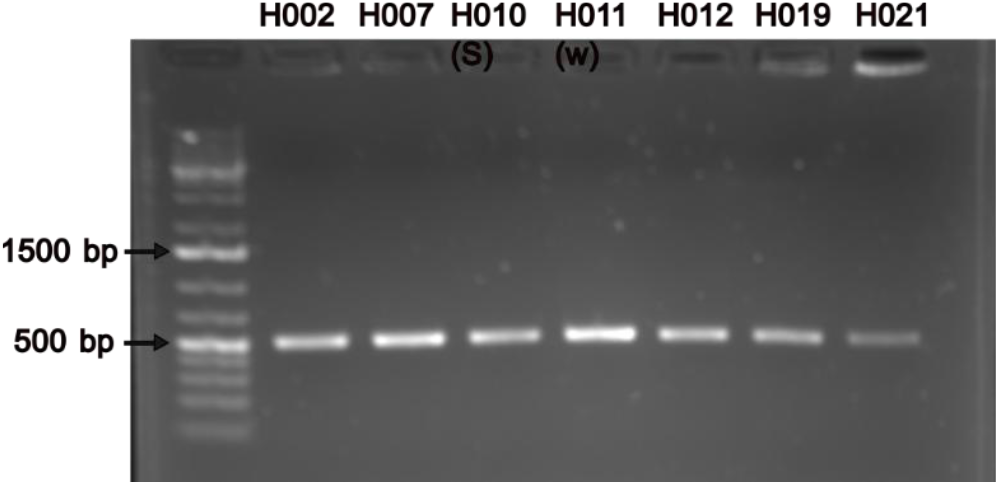
Multiplex PCR to distinguish between *Staphylococcus capitis* and *Staphylococcus caprae*. The *nuc* gene of colonies identified to be either *S. capitis* and *S. caprae* through 16S rRNA sequencing were amplified with stain specific oligonucleotides for amplification of a 525 bp fragment *S. capitis* or a 1227 bp fragment for *S. caprae*. The 500 bp and 1500 bp references are labelled on the Thermofisher GeneRuler™ 1KB plus DNA ladder are marked for reference.

**Supplementary Figure 2:**
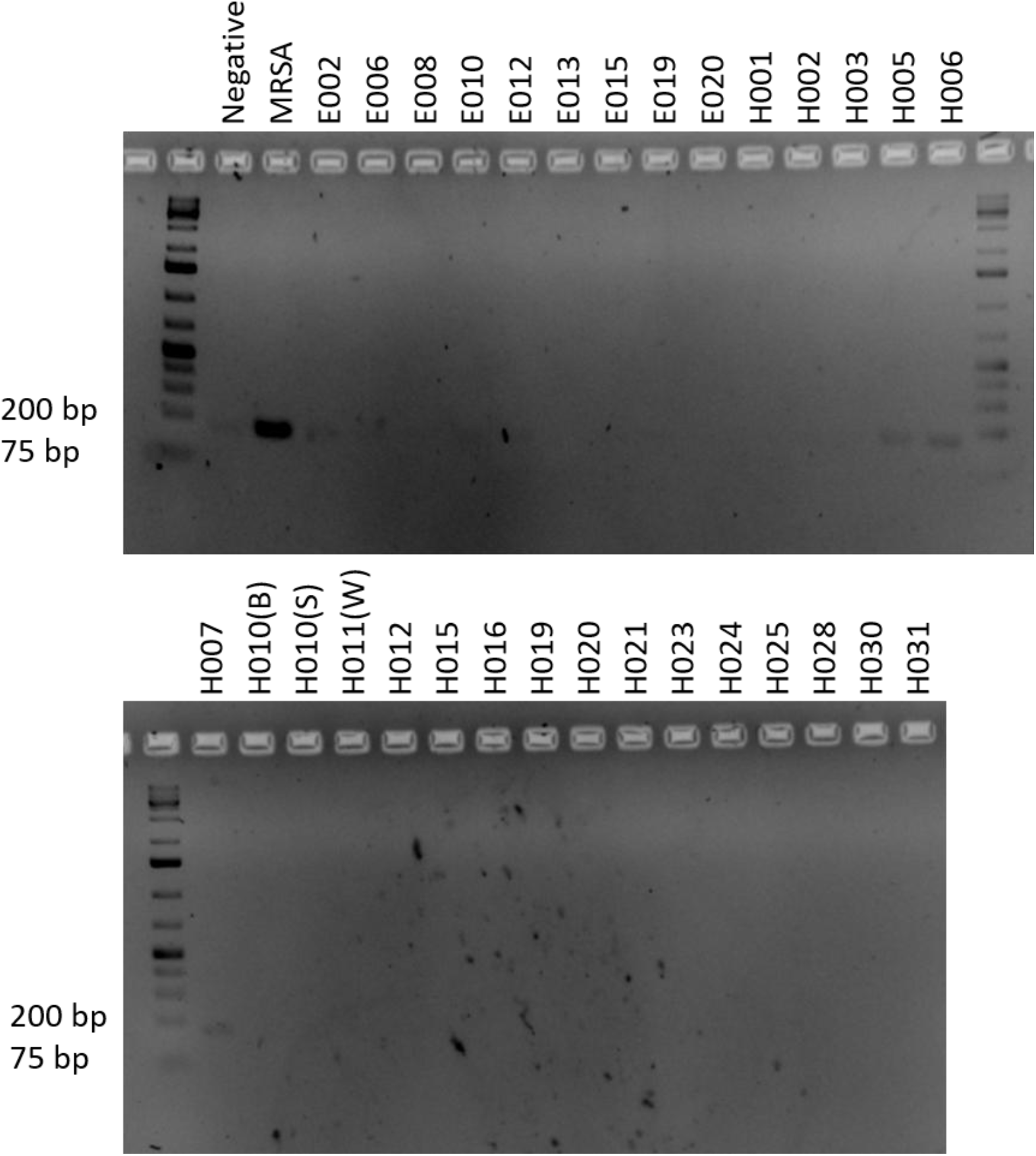
*mecA* PCR to identify staphylococcal strains carrying methicillin resistance. Positive band at 147 bp. PCR products visualised on a 2% agarose TAE gel. The 200 bp and 75 bp references are labelled on the Thermofisher GeneRuler™ 1KB plus DNA ladder are marked for reference.

